# A personalized medicine approach identifies enasidenib as an efficient treatment for IDH2 mutant chondrosarcoma

**DOI:** 10.1101/2023.09.12.557334

**Authors:** Verónica Rey, Juan Tornín, Juan Jose Alba-Linares, Cristina Robledo, Dzohara Murillo, Aida Rodríguez, Borja Gallego, Carmen Huergo, Alejandro Braña, Aurora Astudillo, Dominique Heymann, Agustín F. Fernández, Mario F. Fraga, Javier Alonso, René Rodríguez

## Abstract

**Background:** Sarcomas represent an extensive group of malignant diseases affecting mesodermal tissues. Among sarcomas, the clinical management of chondrosarcomas remains a complex challenge, as high-grade tumors do not respond to current therapies. Mutations in the *isocitrate dehydrogenase* (*IDH*) 1 and 2 genes are among the most common mutations detected in chondrosarcomas and may represent a therapeutic opportunity. The presence of mutated IDH (mIDH) enzymes results in the accumulation of the oncometabolite 2-HG leading to molecular alterations that contribute to drive tumor growth.

**Methods:** We developed a personalized medicine strategy based on the targeted NGS/Sanger sequencing of sarcoma samples (n=6) and the use of matched patient-derived cell lines as a drug-testing platform. The anti-tumor potential of IDH mutations found in two chondrosarcoma cases was analyzed in vitro, in vivo and molecularly (transcriptomic and DNA methylation analyses).

**Findings:** We treated several chondrosarcoma models with specific mIDH1/2 inhibitors. Among these treatments, only the mIDH2 inhibitor enasidenib was able to decrease 2-HG levels and efficiently reduce the viability of mIDH2 chondrosarcoma cells. Importantly, oral administration of enasidenib in xenografted mice resulted in a complete abrogation of tumor growth. Enasidenib induced a profound remodeling of the transcriptomic landscape not associated to changes in the 5mC methylation levels and its anti-tumor effects were associated with the repression of proliferative pathways such as those controlled by E2F factors.

**Interpretation:** Overall, this work provides the first preclinical evidence for the use of enasidenib to treat mIDH2 chondrosarcomas.

**Funding:** Spanish Research Agency (grants PID2019-106666RB-I00; PI20CIII/00020; DTS18CIII/00005; CB16/12/00390; CB06/07/1009; CB19/07/00057).

**RESEARCH IN CONTEXT:** *Evidence before this study:* Sarcomas represent an extensive group of malignant diseases affecting mesodermal tissues. The genomic nature of most sarcoma subtypes, displaying high inter- and intra-tumor heterogeneity with few recurrent driver mutations in a small portion of patients, makes these tumors especially indicated for personalized treatment approaches. For optimal development of these personalized protocols and a more efficient translation to the clinic, it is necessary to create patient-derived models suitable for testing the efficiency of candidate therapies. These strategies might be especially indicated for chondrosarcomas, a subtype of bone sarcoma that is inherently resistant to current therapies.

*Added value of this study:* To develop a personalized medicine strategy for sarcomas we have applied targeted sequencing protocols to detect druggable mutations in a collection of sarcomas cases with available patient-derived models. Among those potential druggable alterations detected in patient samples and avatar cell lines, we found IDH mutations in two chondrosarcomas. The presence of mutated IDH enzymes results in the accumulation of the oncometabolite 2-HG which contributes to driving tumor growth. In vitro and in vivo experiments evidenced the anti-tumor potential of the IDH mutant inhibitor enasidenib for the treatment of IDH2 mutant chondrosarcomas. Our transcriptomic and epigenomic analyses show that the mechanism of action of this drug is associated with the repression of proliferative pathways rather than with the promotion of tumor differentiation.

*Implications of all the available evidence:* This study suggests that enasidenib may represent an efficient therapeutic alternative for mutant IDH2 chondrosarcomas. This anti-proliferative mechanism of action of this drug may be especially relevant in dedifferentiated chondrosarcomas where reversal of this phenotype is not possible. In addition, this work provides support for the use of sarcoma patient-derived lines as avatar models capable of predicting (pre)-clinical responses in personalized medicine strategies.

## INTRODUCTION

The rapid development of Next-Generation Sequencing (NSG) and its implementation in the analysis of molecular alterations in human disease foresee a paradigm shift in the way that cancer patients will be treated in the near future ^1^. It is expected that oncological treatments will be based on the use of personalized strategies for each patient that combine the molecular information gained through NGS with functional drug screenings using patient-derived models to find the most efficient therapy for each case ^2, 3^. These precision medicine approaches will especially benefit those types of tumors that show a high rate of inter- and intra-tumor heterogeneity. This is the case for most types of sarcomas, which are characterized by the presence of few recurrent driver mutations in a small portion of patients ^4, 5^. In sarcomas, cytotoxic drugs (doxorubicin, ifosfamide, cisplatin, etc) and/or radiotherapy remain the cornerstone of systemic treatment and targeted therapies are restricted to the use of tyrosine kinase inhibitors in gastrointestinal stromal tumors harboring mutations in KIT and/or PDGFRA ^6^.

Among sarcomas, chondrosarcomas stand out as the subtype with fewer therapeutic options available. These types of cartilage-forming bone sarcomas are inherently resistant to conventional chemo and radiotherapy and nowadays there are no effective treatments available for metastatic or inoperable tumors ^7–9^. Most common genomic alterations described for chondrosarcoma include mutations in *COL2A1*, *TP53* and specially isocitrate dehydrogenase 1 (*IDH1*) or *IDH2* genes, which has been found in more than 50% of the cases ^10–12^. Mutations in *IDH1/2* genes are also common and play key roles in enchondromas, Ollier disease and Maffucci syndrome, which are benign lesions that may progress to chondrosarcoma ^13, 14^. Also in support of a driver role of mutant *IDH1/2* genes in the origin of chondrosarcomas, mutant IDH enzymes were reported to induce an epigenetic dysregulation of genes involved in stem cell-driven differentiation and lineage specification and the expression of mutant *IDH2* in mesenchymal stem cells induced the formation of sarcomas in vivo ^15,16^. Besides chondrosarcomas, mutations in *IDH* genes are also common and play pro-tumor roles in gliomas (up to 80% of the cases) ^17^, acute myeloid leukemia (AML) (∼20%) ^18^, and cholangiocarcinomas (∼20%) ^19^.

IDH enzymes catalyze the conversion of isocitrate to α-ketoglutarate (α-KG) in the tricarboxylic acid cycle. This enzymatic function is altered due to non-synonymous substitutions of arginine residues R132 in IDH1 and R172 or R140 in IDH2. The mutation of these critical amino acids of the catalytic domain results in the acquisition of a new enzymatic activity in which IDH mutant enzymes catalyze the conversion of α-KG into the oncometabolite d-2-hydroxyglutarate (D2HG) ^20^. The pathological accumulation of D2HG results in the inhibition of α-KG-dependent dioxygenases such as methyl cytosine hydroxylases, and the histone demethylases of the TET family. Inhibition of these enzymes promotes DNA and histone hypermethylation, leading to a transcriptional dysregulation that results in the blockage of different cell differentiation pathways and the onset of oncogenic processes ^9, 21^. In recent years, several inhibitors targeting IDH1/2 mutants have been developed ^22, 23^. Some of these compounds have been demonstrated to be effective in pre-clinical studies, mainly in AML and gliomas, and are being tested in clinical trials ^22–24^. Among them, AG-120 (ivosidenib) and AG-221 (enasidenib), which are specific inhibitors against IDH1 and IDH2 mutants respectively, are first-in-class compounds and has been approved by FDA for the treatment of relapsed or refractory AML ^25,26^. IDH anti-mutant treatments may also represent a therapeutic alternative for chondrosarcomas, however, the effect of these compounds in this type of tumors has been barely explored. On one hand, different studies using IDH1 mutant inhibitors reached contradictory conclusions about the ability of this drug to suppress tumorigenic properties in chondrosarcoma cells ^27–29^. In the clinical setting, Ivosidenib showed minimal toxicity and durable disease control in a small cohort of chondrosarcoma patients enrolled in a phase I study ^30^. On the other hand, the effect of IDH2 mutant inhibitors in chondrosarcoma had not been tested so far.

Here we aimed to assay a personalized medicine strategy for sarcomas based on the use of patient-derived primary cell lines as a drug-testing platform. Among those potential druggable alterations found in six sarcoma cases, we found mutation IDH mutations in two chondrosarcomas. In vitro and in vivo experiments evidenced the anti-tumor potential of enasidenib for the treatment of IDH2 mutant chondrosarcomas.

## METHODS

### Cell lines

CDS11 and CDS17 patient-derived lines and T-CDS17 xenograft lines were previously established and described ^31^. CDS20, SARC06, SARC20 and SYN01 lines were also established following the same methodology. L2975 cells were described elsewhere ^32^. The SW1353 cell line was purchased from the ATCC. An overview of patient and tumor characteristics associated with each cell line is given in Table S1. All the cell types were cultured as previously described ^31, 33^ (supplemental information).

### Drugs

Enasidenib (AG-221) was purchased from MedChemExpress (Monmouth Junction, NJ, USA) and ivosidenib (AG-120), vorasidenib (AG-881), metformin and CB-839 were acquired from Selleckchem (Houston, TX, USA). Stocks for in vitro experiments were prepared as 10 mM solutions in sterile DMSO, stored at −80 °C and diluted in culture medium to the final concentration just before use. For in vivo treatments, enasidenib was solubilized in distilled water containing 0.5% methylcellulose (Sigma-Aldrich, St Louis, MO, USA) and 0.2% Tween-80 (Sigma-Aldrich) before administration.

### Next Generation Sequencing

Genomic DNA from matched samples of healthy tissue (blood), tumor tissue and tumor-derived cell lines was extracted using the QIAmp DNA Mini Kit (Qiagen, Hilden, Germany). Subsequent targeted sequencing of 160 cancer-related genes was performed using a QIAseq Human Comprehensive Cancer Panel v2 (Qiagen Inc, Valencia, CA) (Table S2). In addition, the panel includes more than 21 genes in which clinically actionable alterations have been described, as well as 29 genes known to predispose to cancer. Multiplex PCR was performed by amplifying 10 ng of DNA in each of 4 separate PCR reactions to generate the corresponding libraries for next-generation sequencing. All steps of library preparation were performed according to the manufacturer’s protocol (see Supplemental information). For the Human Comprehensive Cancer Panel, up to 12 samples were pooled equimolar and paired-end reads 2×150bp sequencing on an Illumina NextSeq 500 instrument using the 300 cycle Mid Output v2.5 Reagent Kit (Illumina, San Diego, CA, USA) according to Illumina guidelines. Read mapping, variant discovery, and functional annotation were performed with the Qiagen GeneRead Variant Calling Pipeline. The identification and clasification of somatic mutations were performed as described in Supplemental Information.

The mutational status of *IDH1*, *IDH2*, *SETD2*, *TSC1* and *TP53* was confirmed by Sanger sequencing as described before ^31^. PCR reactions for Sanger sequencing were carried out using the primers detailed in Table S3.

### RNA sequencing

RNA samples were extracted from triplicated cultures of control and enasidenib treated cell lines as previously described ^34^. cDNA libraries were prepared using the TruSeq RNA Library Preparation Kit (Illumina) and checked for quality and quantified using a TapeStation (Agilent Tecnologies). Paired-end sequencing was performed on an Illumina Novaseq 6000 (Illumina) using 100-base reads. RNA-seq data processing, differential gene expression analyses, gene annotation and gene set enrichment analyses were performed as described in Supplemental Information.

#### Microarray analysis of chondrosarcoma samples

Microarray expression data (Affymetrix Human Gene 2.0 ST arrays) from Nicolle et al. (2019) were analyzed to determine whether enasidenib treatment can alter transcriptome profiles associated with chondrosarcoma expression subtypes. Log expression values from microarray and mRNA-seq experiments were independently normalized to the [0,1] range to reduce bias due to the different expression technologies considered, and samples were hierarchically clustered using Euclidean distances.

### Genome-wide DNA methylation analysis

Genomic DNA was extracted from triplicated cultures of control and enasidenib treated cell lines as described above. The EZ-96 DNA Methylation Kit (Zymo Research) was used to perform the bisulphite conversion. The processed DNA samples were then hybridised to Infinium MethylationEPIC BeadChips following the Illumina Infinium HD Methylation Assay protocol. Data processing, differential methylation analyses, probe annotation and pathway enrichment analyses were performed as described in Supplemental Information.

#### Integration of methylation array data from cartilage samples

Appropriate publicly available methylation datasets were considered to characterize the methylation landscape of healthy human cartilage samples (ArrayExpress accession codes: E-GEOD-63106, E-GEOD-73626). HumanMethylation450 BeadChip arrays from cartilage samples were integrated with MethylationEPIC BeadChip arrays to extract common CpGs between the two technologies (399,453 probes). All cartilage samples passed the quality controls applied to cell line samples, including validation of self-reported sex.

### Cell viability assays

Cell viability of cell lines in the presence of increasing concentrations of different drugs was assayed using the cell proliferation reagent WST-1 (Roche, Mannheim, Germany) after 72 h-treatments as previously reported ^35, 36^. IC_50_ for each treatment was determined by non-linear regression using GraphPad Prism 9.0.1 software (Graphpad Software Inc., San Diego, CA, USA). Alternatively, the cytotoxic potential of the assayed treatments was assayed in colony formation unit (CFU) assays as described previously ^37^. The induction of apoptosis was assayed using the Alexa Fluor 488 Annexin V / Dead Cell Apoptosis Kit (Life Technologies, San Francisco, USA) following the manufacturer’s instructions. All cell viability assays were performed in the presence of fetal bovine serum.

### D-2-Hydroxygluturate measurement

D2HG levels were measured in adherent cultures of 1 x 10^7^ chondrosarcoma cells treated with increasing concentrations of Enasidenib for 48 hours using the D2HG colorimetric Assay Kit (Abcam, Cambridge, UK) following the manufactureŕs recommendations.

### Tumorsphere formation assay

Tumorsphere formation protocol and the analysis of the effects of drugs on tumorsphere formation were performed as previously described ^38, 39^.

### Immunofluorescence staining

The formation of DNA damage-induced γ-H2AX foci was analyzed by Immunofluorescence staining using a mouse monoclonal anti-phospho-Histone H2A.X (Ser139) (Sigma) as described before ^40^. Quantification of dots per nuclei was performed in 100-120 randomly selected cells using Image J software (National Institute of Health, Bethesda, MD).

### Chondrogenic differentiation assay

To induce chondrogenic differentiation, 2.5 x 105 cells were placed in a 15 ml polypropylene tube (Corning, TX, USA) and centrifuged at 1.000 rpm for 3 min. The pellet was cultured in 500 µl of either standard culture medium or ready-to-use complete chondrogenic differentiation medium StemPro chondrogenesis differentiation kit, Gibco-Thermo Fisher, Waltham, MA, USA) following the manufactureŕs recommendations. The medium was carefully removed to avoid pellet resuspension and replaced every 3 days with or without drugs for 21 days. Then, pellets were washed in PBS, fixed at 4°C with 4% formaldehyde and embedded in HistoGel (Epredia, Kalamazoo, MI, USA) firstly and in paraffin afterwards processed for histological analysis (H&E and PAS-alcian blue stainings) as described below. Slides were scanned with the digital scanner Motic EasyScan One (Motic, Xiamen, China) and the percentage of PAS-alcian positive areas relative to the total area was quantified using the ImageJ 2.1.0 software (National Institutes of Health, Bethesda, MD, USA)″

### Treatment of tumor xenografts

Athymic nude-Foxn1nu mice (Envigo, Barcelona, Spain) of 6 weeks old were inoculated subcutaneously (s.c.) with 5 × 10^5^ cells mixed 1:1 with BD Matrigel Matrix High Concentration (BD Biosciences, Erembodegem, Belgium) previously diluted 1:1 in culture medium. Once tumors reached approximately 100 mm3, the mice were randomly assigned (n = 6 per group) to receive oral gavage doses of enasidenib (35 mg/kg per dose) or vehicle (0.5% methylcellulose and 0.2% Tween-80) twice a day (b.i.d.) for 21 days. Tumor size was measured using a caliper 3 times a week and tumor volume was determined using the equation (D × d2)/6 × 3.14, where D is the maximum diameter, and d is the minimum diameter. Relative tumor volume (RTV) for every xenograft was calculated as follows: RTV = tumor volume at the day of measurement (Vt)−tumor volume at the beginning of the treatment (Vo). At the end of the treatment, mice were sacrificed by cervical dislocation and tumors were extracted, fixed in 4% formaldehyde and processed for histological analysis. This was a nonblinded study.

### Histological analysis

Xenograft tumor samples were fixed in 4% formaldehyde, embedded in paraffin, cut into 4-µm sections, and stained with hematoxylin and eosin (H&E) and PAS-alcian blue staining as previously described ^41, 42^. Immunohistochemical analyses were performed in an automatic workstation (Dako Autostainer Plus) with anti-Ki67 (Clone MIB-1 Dako # JR626, Prediluted), using the Dako EnVision Flex + Visualization System (Dako Autostainer) ^43^. Quantification of Ki67 staining was performed by counting the number of positive cells per 10 high-power fields (40×). To quantify PAS-alcian staining, slides were scanned with the digital scanner Motic EasyScan One (Motic, Xiamen, China) and the percentage of PAS-alcian positive areas relative to the total area was quantified using the ImageJ 2.1.0 software (National Institutes of Health, Bethesda, USA) in three random images (x200) per condition.

### Prediction of protein conformation

Wild-type IDH1 and 2 structures were obtained in the AlphaFold Protein Structure Database (https://www.alphafold.ebi.ac.uk/). The in-silico mutation impact on the protein structure was determined via the Dynamut server (https://www.biosig.lab.uq.edu.au/dynamut/prediction) ^44^. PDB files were visualized using UCSF ChimeraX program (https://www.cgl.ucsf.edu/chimerax/).

### Statistical Analysis

Statistical analysis was performed using GraphPad Prism 9.0.1 software. Unless otherwise indicated, all data are presented as the mean (±standard deviation or SEM as indicated) of at least three independent experiments. A two-sided Student’s t-test was performed to determine the statistical significance between groups. Multiple comparisons of the data were performed using one-way ANOVA and Tukey’s test, p ≤ 0.05 values were considered statistically significant.

### ethics statement

All experimental protocols involving human samples were conducted following the institutional review board guidelines and in compliance with the WMA Declaration of Helsinki and were approved by the Institutional Ethics Committee of the Principado de Asturias (ref. 255/19). All animal research protocols were approved by the Animal Research Ethical Committee of the University of Oviedo (ref. PROAE 34-2019).

### Data deposition

Raw data files obtained from RNA seq and DNA methylation analysis have been deposited at the GEO-NCBI repository with SuperSeries reference GSE235701 (https://www.ncbi.nlm.nih.gov/geo/query/acc.cgi?acc=GSE235605 and https://www.ncbi.nlm.nih.gov/geo/query/acc.cgi?acc=GSE235697).

## RESULTS

### Identification of targetable mutations in sarcoma patients

We aimed to find actionable mutations in sarcoma cases with available matched patient-derived cell lines. In this analysis, we included tumors and primary cell lines (CDS11 and CDS17) conventional chondrosarcoma (CDS11) and an dedifferentiated chondrosarcoma (CDS17), as well as several cells lines established from CDS17-developed xenografts (T-CDS17#1, #3, and #4), which have been previously characterized ^31^. In addition, we analyzed two cases of undifferentiated pleomorphic sarcomas (SARC06 and SARC20), one synovial sarcoma (SYN01) and a extraskeletal myxoid chondrosarcoma (CDS20) with newly developed cell lines (Table S1). First, we performed NGS targeted sequencing of 160 cancer-related genes (Table S2) in normal (non-tumoral) tissue, tumor samples and patient-derived models of SARC06, SAR20, SYN01 and CDS17 patients. The average nucleotide coverage in DNAseq studies was > 2500 reads in all samples analyzed (range 2488-5340 reads), and only the variants presenting more than 100 reads were selected for further analyses. Data from NGS analysis of the tumor and cell line samples were compared to that of normal tissue DNA to exclude germline alterations. To select those somatic mutations with potential driver roles in each case, we filter the results to select variants with non-synonymous effects on coded proteins which presented variant allele frequencies >0.30 in tumor and cell line samples and maximum allele frequencies <0.01 in population databases (dbSNP, ExAC, ESP, and 1000 Genomes) (Figure 1A). Using these criteria, we found 3 somatic mutations that were shared by both tumor and cell line samples: *SETD2* (p.E2089*) and *TP53* (p.V173L) in SARC06 and *IDH2* (p.R172G) in CDS17 / T-CDS17#1. In addition, we found mutations in *TSC1* (p.SS331R) and *TP53* (p.S215R) that appears de novo in the cell lines SARC20 and CDS17 / T-CDS17#1 respectively, thus reflecting a certain level of genetic drift of tumor cells during their adaptation to in vitro growth conditions. Finally, we did not find relevant mutations in the SYN01 case. All detected mutations in NGS panels were confirmed by Sanger sequencing (Figure 1B and Figure S1). In addition, Sanger sequencing analysis also confirmed the presence of the *IDH2* (p.R172G) mutation in other cell lines (T-CDS17#3 and T-CDS17#4) derived from CDS17-generated xenografts (Figure 1A and Figure S2). As previously seen, we also detected a heterozygous mutation in *IDH1* (p.R132L) in CDS11 samples (Figure 1A-B) ^31^. Finally, we did not find *IDH* mutations in the extraskeletal myxoid chondrosarcoma CDS20 case (Figure 1A and Figure S2). Most common mutations reported in the R172 residue of IDH2 in chondrosarcoma include variants p.R172S (50%), p.R172G (30%), p.R172W (10%) and p.R172T (10%) ^45^. On the other hand, reported variants in the R132 residue of IDH1 in chondrosarcomas include p.R132C (44%), p.R132G (17%) and p.R132L (11%) among others ^45^. As depicted in Figure 1C-D, the R172 arginine in IDH2 and R132 in IDH1 are located in the isocitrate-binding pocket where these and other residues activated the catalytic conversion of isocitrate to α-KG ^46^. The presence of R172S / G / or W variants in IDH2 or R132C / G / or L variants in IDH1 induces conformational changes in isocitrate-binding pocket of these enzymes that may lead to the formation of D2HG (Figure 1C-D).

**Figure 1.**
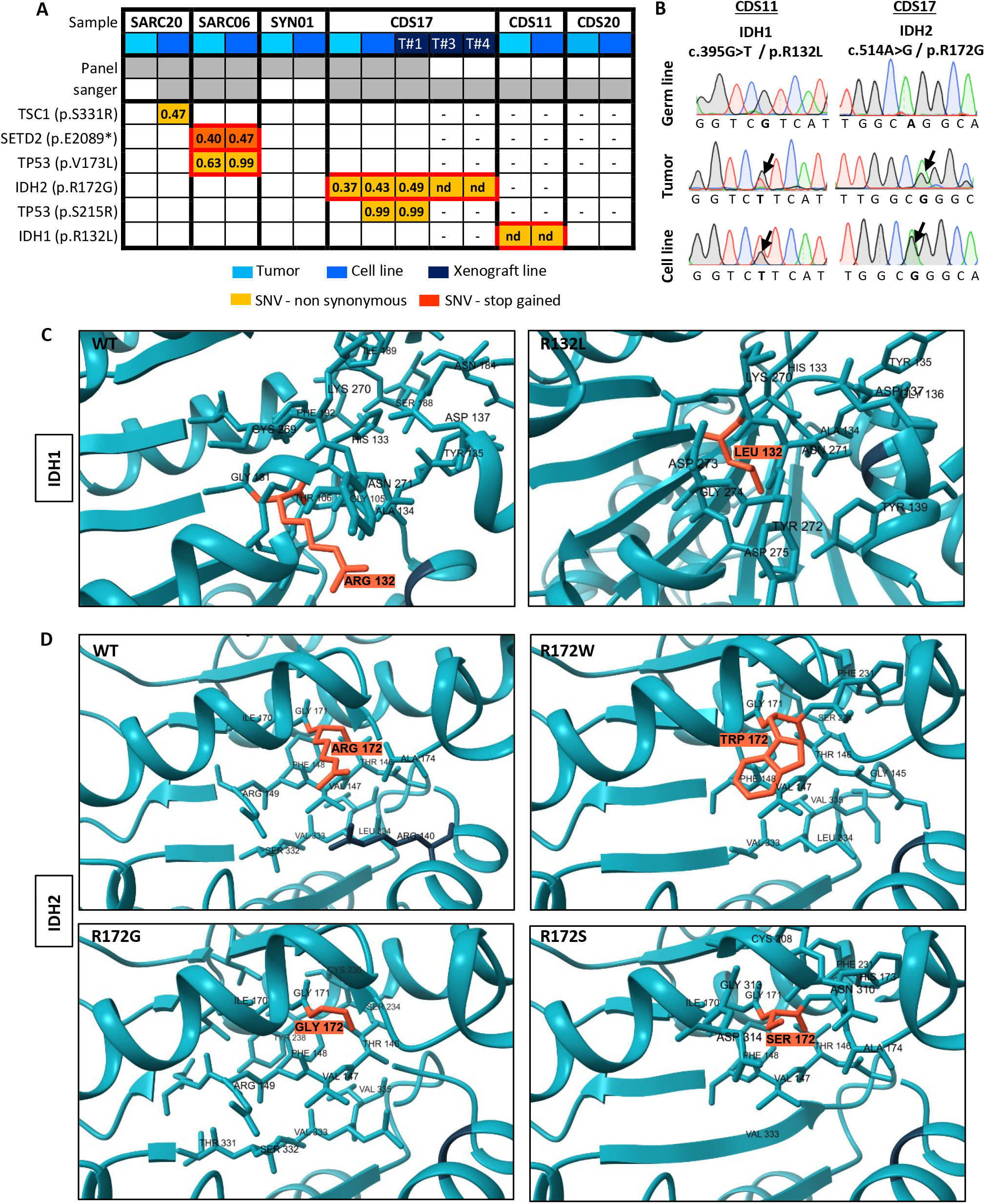
Mutations found in patient-derived sarcoma models. (A) Summary of somatic mutations found by NSG-panel and/or Sanger sequencing in tumor samples and/or primary cell lines derived from six sarcoma patients. Variant allele frequency for each mutation detected by NSG sequencing is indicated. (B) Sanger sequencing validation of IDH mutations found in CDS11 and CDS17 models (Sanger validation of the rest of mutations is presented in Supp Figs 1 & 2). (C-D) In silico docking modelling of the conformational changes caused by the mutation in IDH1 (C) and IDH2 (D) in the substrate-binding pocket. Residues 132 in IDH1 and 172 in IDH2 are highlighted.

### Targeting IDH mutations in chondrosarcoma

Given the prominent role of IDH mutations in chondrosarcomas and the availability of drugs specifically directed against these mutations, we focused on studying the effect of targeting mutant IDH enzymes in the chondrosarcoma models. In these analyses, we also included two previously reported chondrosarcoma cell lines with IDH2 mutations: SW1353 (IDH2-R172S) and L2975 (IDH2-R172W) ^32^ (Table S1). In dose-response cell survival assays, we found that the CDS17 cells and their associated xenograft-derived cell lines T-CDS17#1, #2 and #3, carrying the IDH2-R172G variant, were sensitive to low µM concentrations of IDH2mut inhibitor enasidenib (IC_50_ between 16.65 and 22.65 µM.) (Figure 2A-B). In these experiments SW1353 cells (IDH2-R172S; IC_50_= 49.47 µM) displayed an intermediate sensitivity, whereas CDS11 (IDH1-R132L; IC_50_= 63.53) and L2975 cells (IDH2-R172W; IC_50_= 73.53 µM) were partially resistant to enasidenib. Finally, extraskeletal myxoid chondrosarcoma CDS20 cells (IDH-wt; IC_50_>100 µM) we fully resistant to this drug (Figure 2A-B). On the other hand, when treated with the IDH1mut specific inhibitor ivosidenib (Figure 2C) or the dual inhibitor of both IDH1 and IDH2 mutants vorasidenib (AG-881) (Figure 2D), none of the assayed lines showed a significant response to these drugs.

**Figure 2.**
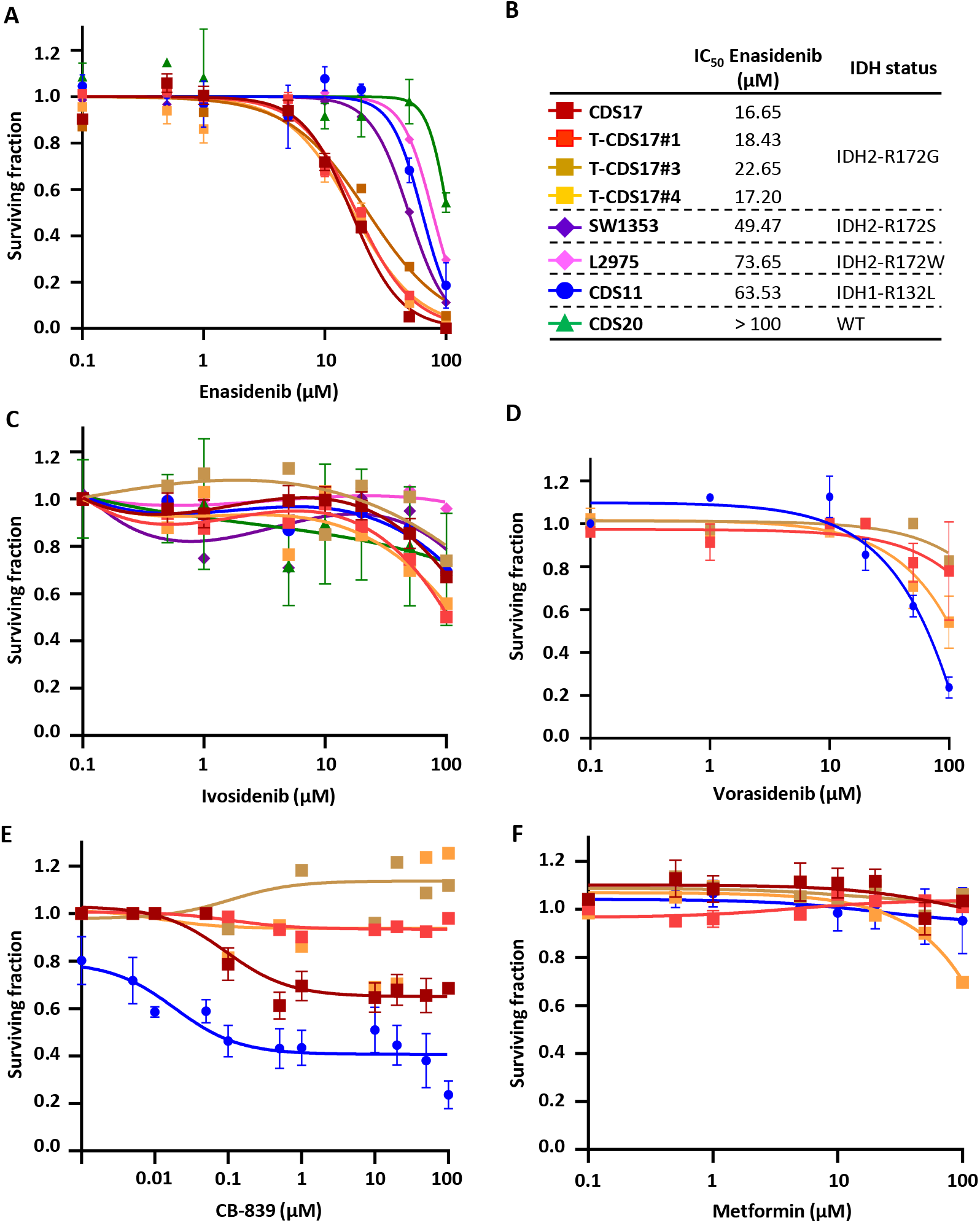
Dose-dependent effect of several inhibitory compounds on chondrosarcoma cell lines. (A-F) Cell viability (WST-1 assays) measured after the treatment of wild-type and mutant IDH chondrosarcoma cell lines with increasing concentrations of enasidenib (A), ivosidenib (C), vorasidenib (D), CB-839 (E) and metformin (F) for 72 h. IDH mutation status of chondrosarcoma cell lines and IC5_50_ values for enasidenib treatments are shown (B).

Another type of anti-IDHmut therapy tested in chondrosarcomas is based on the inhibition of pathways that lead to the synthesis of α-KG, the substrate that IDH1 and IDH2 mutant enzymes use to produce D2HG. Therefore, inhibitors of glutaminolysis such as metformin and CB-839 have been tested in different chondrosarcoma cell lines ^47^. In these experiments, only CB-839 was able to partially reduce cell viability in the IDH1 mutant cell line CDS11 but not in the CDS17/T-CDS17 IDH2 mutant models (Figure 2E), while metformin treatment did not affect cell viability in any of these cell lines (Figure 2F).

Further exploring the antiproliferative effects of enasidenib in IDH mutant models, we performed colony-forming assays with T-CDS17#1, SW1353 and L2975 IDH2 mutant cells following the treatment with increasing concentrations of enasidenib. Similar to our results in cell viability assays, we found that T-CDS17#1 and SW1353 lines were the most sensitive lines, whereas L2975 cells displayed a resistant phenotype (Figure 3A-B). The cytotoxic effect of enasidenib was partially mediated by the induction of apoptosis, as evidenced by the time-dependent accumulation of the T-CDS17#4 cells staining positive for annexin V / propidium iodide after drug treatment (figure S3A-B). In any case, the induction of apoptosis was not associated with the induction of DNA damage (figure S3C-D).

**Figure 3.**
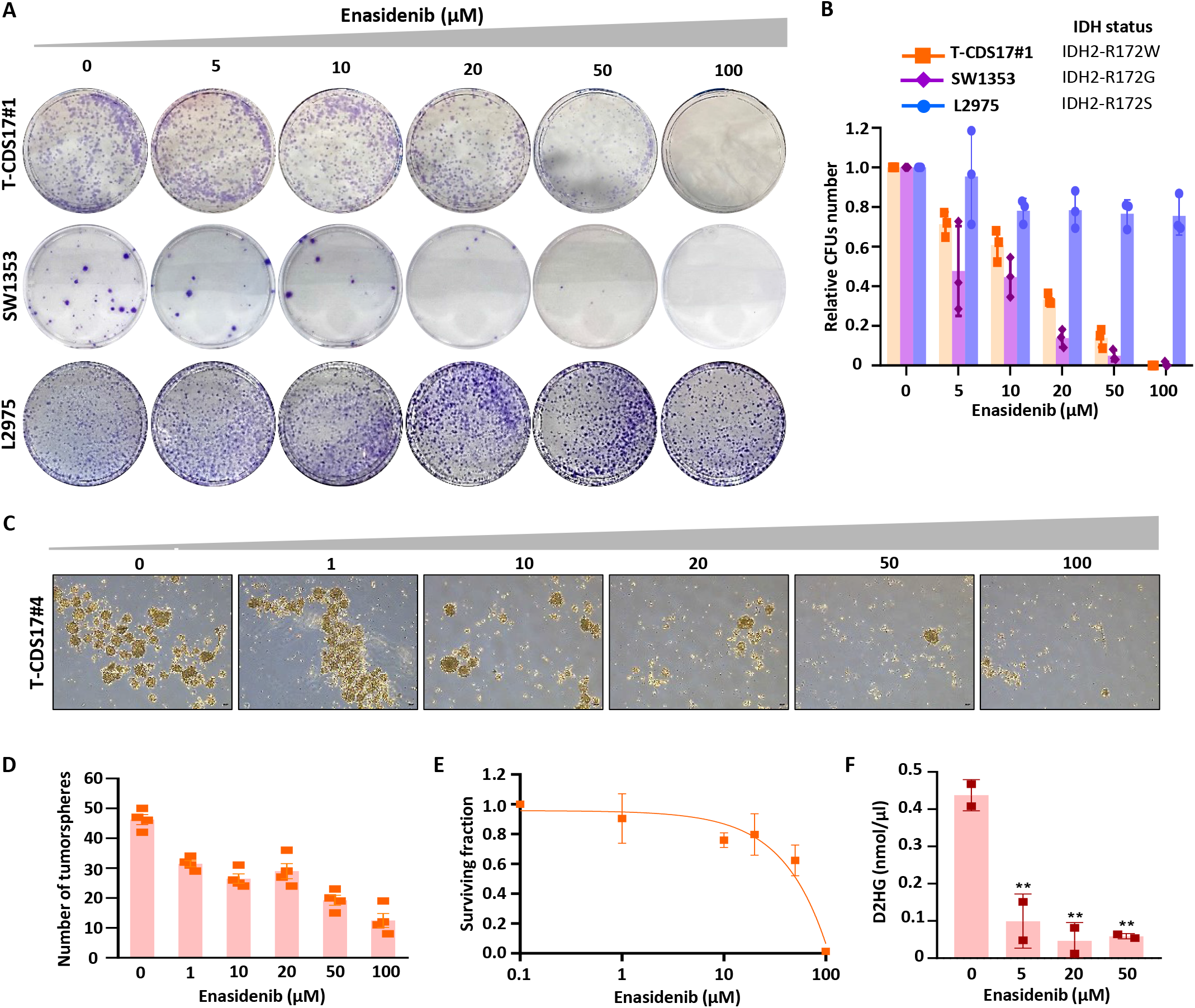
Effect of enasidenib in IDH2 mutated chondrosarcoma cell lines. (A-B). Colony formation unit (CFU) assay of T-CDS17#1, SW1353 and L2975 cells treated with increasing concentrations of enasidenib for 24 h and left to form CFUs for 10 days. Representative pictures (A) and quantification (B) of a CFU assays for each cell line are shown. (C-E) CSC-enriched tumorspheres of T-CDS17#4 cells were treated with increasing concentrations of enasidenib for 96 h. Representative pictures of the spheres cultures (C), quantification of the number of spheres (D) and cell viability (WST-1 assay) (E) of spheres at the end of the treatment are shown. Scale bar = 100 μm. (F) Intracellular D2HG levels were determined in T-CDS17#1 cells treated or not with the indicated concentrations of enasidenib for 48 h. Error bars represent the standard deviation of two independent experiments for panel F and three independent experiments for the rest of the panels. Asterisks indicate statistically significant differences (**:p<0.01 one-way ANOVA).

Cell lines of the CDS17 model were able to grow as 3D tumorspheres with enhanced cancer stem cell (CSC)-related properties, as previously shown ^31^. To test whether enasidenib may also target these 3D models, we treated tumorsphere cultures of T-CDS17#1 cells with increasing concentrations of the inhibitor for 96 hours. Following this treatment, tumorspheres displayed a disrupted and irregular morphology and their number and viability dramatically decreased in a dose-dependent fashion (Figure 3C-E).

According to the robust effect of enasidenib in the CDS17 model, we confirmed that the treatment of T-CDS17#1 cells with this inhibitor produced a strong dose-response reduction in the intracellular levels of D2HG (Figure 3F).

### DNA methylation and gene expression profiling of enasidenib-treated chondrosarcoma cells

The therapeutic activity of enasidenib and other IDH mutant inhibitors in AML and glioma was associated with the recovery of the D2HG-inhibited activity of αKG-dependent dioxygenases and the subsequent restoration of normal DNA methylation levels ^24^. To analyze the effect of enasidenib on the methylome of chondrosarcoma, we first performed exploratory analyses to describe the global landscape of DNA methylation alterations in control and enasidenib-treated T-CDS17#1 cells, along with healthy cartilage samples. As expected, T-CDS17#1 cells underwent a total remodeling of DNA methylation profiles in comparison with healthy cartilage (Figure 4A). However, T-CDS17#1 cells treated with enasidenib did not show any noticeable differences in global methylation values compared to control cells (Figure 4A-B). Consequently, we performed differential analyses to identify genome-wide differentially methylated positions (DMPs, FDR<0.05, |Δβ|>20%) across the different groups (Figure 4C). Using empirical Bayes moderated t-tests, we discovered abundant DNA methylation alterations between control T-CDS17#1 cells and cartilage samples (128,826 CON vs CART DMPs), and also when compared to enasidenib-treated cells (129,083 ENA vs CART DMPs). Despite this profound remodeling, we detected only 1 hypomethylated DMP between enasidenib-treated cells and their control counterparts, so the treatment did not induce representative alterations at the methylome level. To verify the low impact of enasidenib treatment on DNA methylation, we evaluated the consistency between the DMPs identified in the CON vs CART and ENA vs CART comparisons, showing a high concordance (126,762 COMMON DMPs). Furthermore, the direction of DNA methylation alterations was well balanced in the comparisons CON vs CART (69,765 hypermethylated DMPs, 59,061 hypomethylated DMPs), ENA vs CART (69,419 hypermethylated DMPs, 59,664 hypomethylated DMPs) and COMMON (68,531 hypermethylated DMPs, 58,231 hypomethylated DMPs).

**Figure 4.**
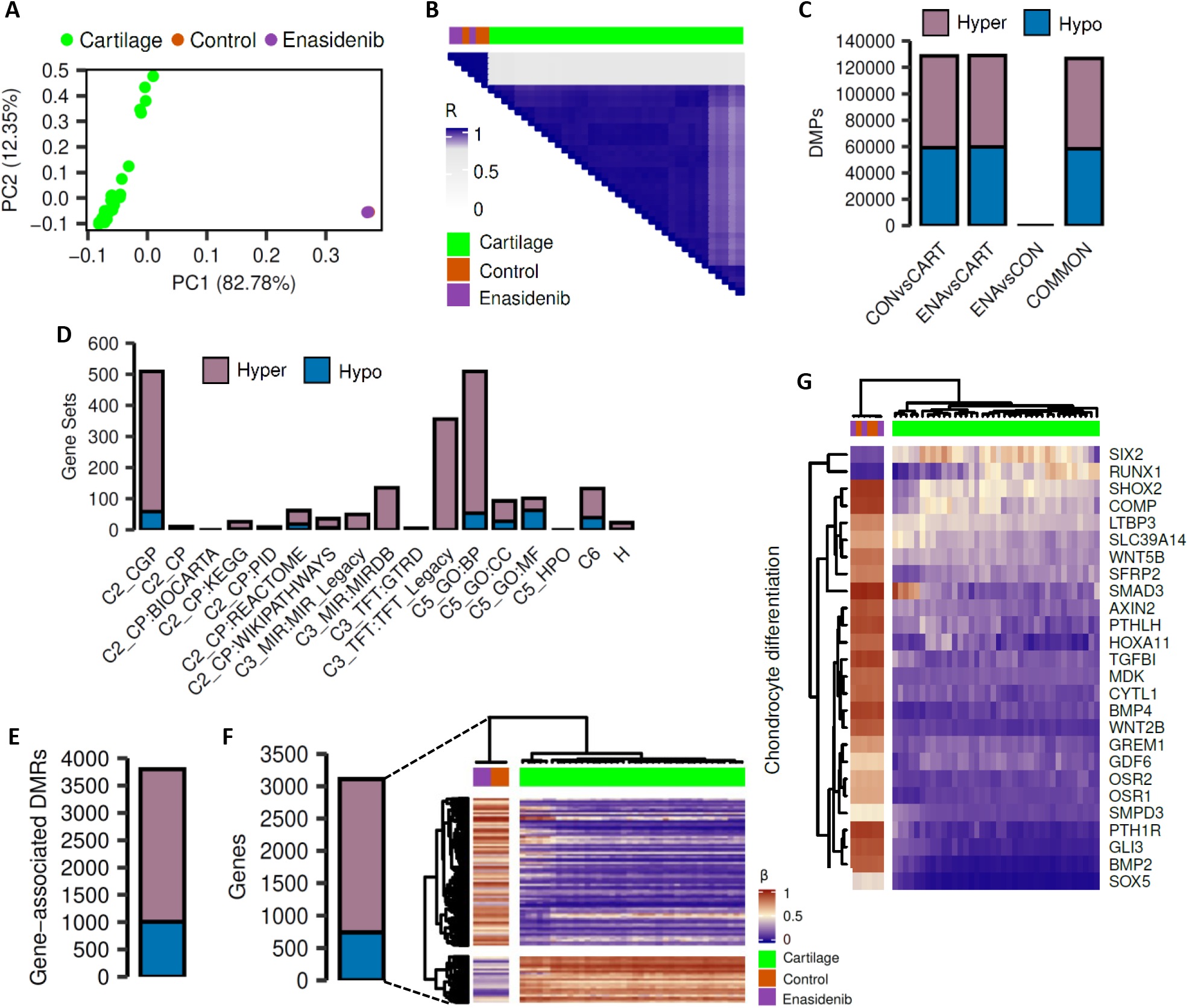
Effect of enasidenib on the methylome of IDH2-mutant chondrosarcoma cells. T-CDS-17#1 cells were treated in triplicates with DMSO (control; CON) or 20 µM enasidenib (ENA) for 48 h prior to be processed for DNA methylation analysis. Public-available methylation datasets of healthy human cartilage samples (CART) were included in these analyses. (A) Scatter plot showing the PCA of the samples according to the beta methylation values at the top 50,000 most variable CpG sites. (B) Heatmap plot showing Pearson’s correlations between beta methylation values of all samples. (C) Barplots depicting the number of hyper- and hypomethylated DMPs (FDR<0.05, |Δβ|>20%) found in CON vs CART, ENA vs CART and ENA vs CON comparisons. COMMON bar includes those hyper- and hypo-DMPs that are in both CON vs CART and ENA vs CART comparisons. (D) Barplots showing the number of significantly enriched pathways found using hyper- and hypomethylated common DMPs for different MSigDB gene sets. (E) Barplot depicting the number of hyper- and hypomethylated DMRs (FDR < 0.05, |Δβ|>20%) found in gene regions using common DMPs. (F) On the left, a barplot showing the number of hyper- and hypomethylated genes. On the right, a heatmap plot showing the beta methylation values of the aforementioned differentially methylated genes. (G) Heatmap plot depicting the beta methylation values of those differentially methylated genes that belong to the GO BP chondrocyte differentiation pathway.

When we studied the functionality of the COMMON DMPs identified, we found that the majority of the enriched pathways were associated with hypermethylation alterations across the different molecular databases from the MSigDB (Figure 4D). Consequently, although the number of methylation alterations is directionally balanced, only the methylation gain is functionally relevant to explain the epigenetic differences between healthy cartilage and T-CDS17#1 cells. Next, to further increase our power to detect functional DNA methylation alterations, we performed analysis at the regional level to define differentially methylated regions (DMRs, FDR<0.05, |Δβ|>20%). We then detected 2789 hypermethylated DMRs and 1009 hypomethylated DMRs (Figure 4E), involving a total of 2367 hypermethylated and 741 hypomethylated genes (Figure 4F). Therefore, in comparison with healthy cartilage, the IDH2 mutant T-CDS17#1 cell line showed a high remodeling of their DNA methylation landscape, with hypermethylated DMPs being more functionally relevant. Among the most significantly altered pathways between healthy cartilage and the T-CDS17#1 cell line, intense hypermethylation of genes involved in chondrocyte differentiation stands out (Figure 4G). In any case, this hypermethylated status of genes was not affected by the enasidenib treatment of tumor cells (Figure 4F-G).

In sharp contrast with this finding, RNAseq analyses of these samples showed that enasidenib induced a profound remodeling of the transcriptomic landscape of T-CDS17#1 cells (Figure 5A). Therefore, we selected differentially expressed genes (DEG) (log_2_ fold change ≤-1 or ≥1 and FDR < 0.01) in treated vs control samples. We found a higher proportion of downregulated (1561) vs. upregulated genes (550) (Figure 5B). Enasidenib treatment resulted in the downregulation of genes like *LIF*, *KIT*, *WNT7B*, *ACVRL1* or *BMP2* and the upregulation of *RSP03*, *IL1RL1*, *GDF15*, *BEX2* or *DDIT4* genes among others (Figure 5C).

**Figure 5.**
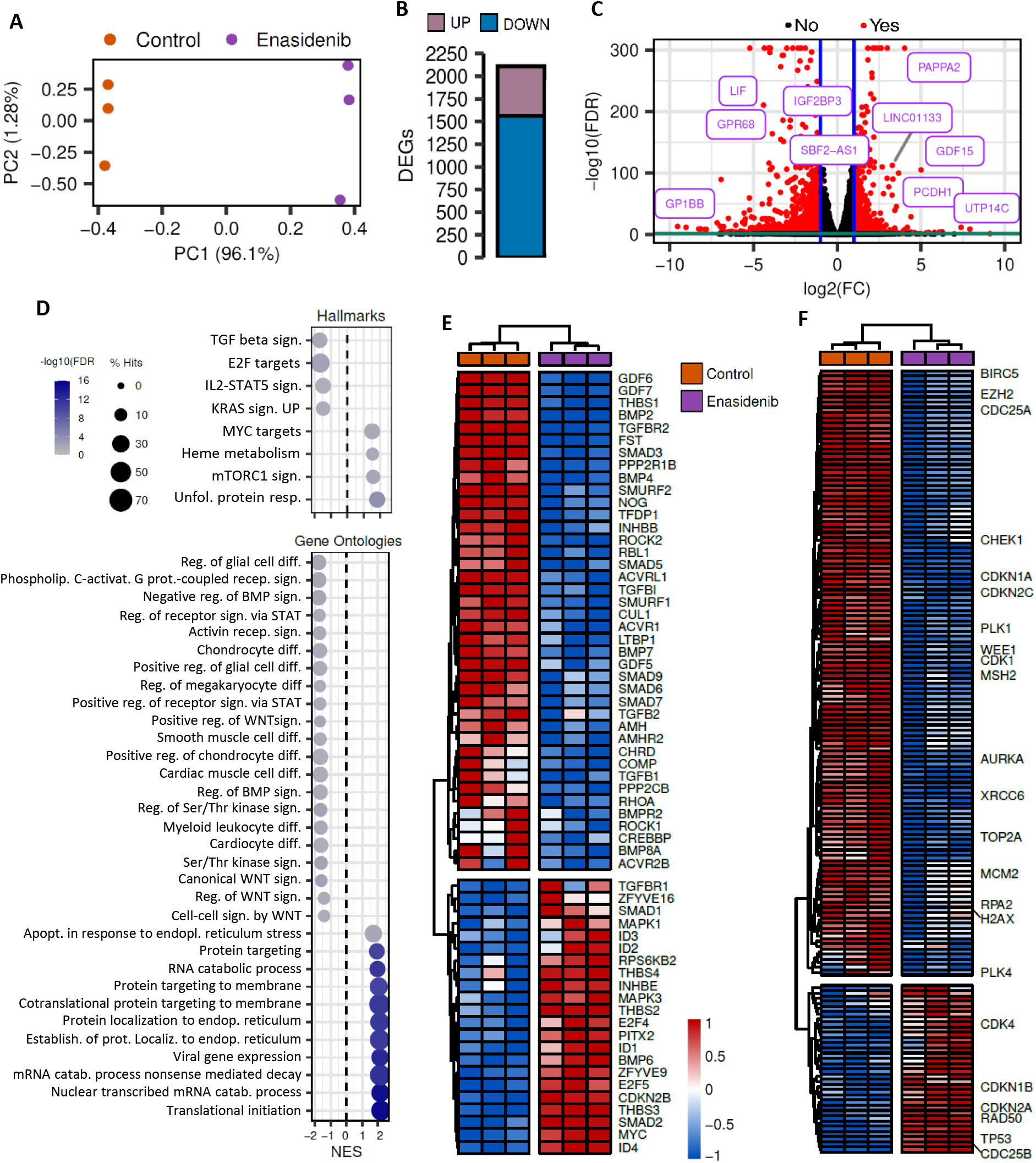
Effect of enasidenib on the transcriptome of IDH2-mutant chondrosarcoma cells. T-CDS-17#1 cells were treated in triplicates with DMSO (control) or 20 µM enasidenib for 48 h prior to be processed for RNA sequencing. (A) Scatter plot showing the PCA of the samples according to the rlog expression values at the top 1,000 most variable genes. (B) Barplot depicting the number of up- and down-regulated DEGs (FDR<0.05, |log2(FC)|>1). (C) Volcano plot showing DEGs (FDR<0.05, |log2(FC)|>1|) in red color. Selected genes displaying highly significant FDRs and/or high fold change are indicated. (D) Bubble plots showing selected significantly enriched pathways (FDR<0.05) of the MSigDB Hallmark (top) and MSigDB GO BP (bottom) collections. Genes appearing in the leading-edge subset are considered positive hits. (E-F) Heatmap plots depicting the expression values of those DEGs (FDR<0.05) of the KEGG TGFβ signalling pathway (E)and the Hallmark E2F targets (F).

GSEA analysis of DEGs revealed the upregulation of relevant pathways such as the unfolded protein response, mTORC1 signaling or the regulation of MYC targets. On the other hand, signaling pathways such as TGFβ signaling, E2F targets, IL2-STAT5 signaling, KRAS signaling or the positive regulation of STAT and WNT signaling were found downregulated (Figure 5D). Among these pathways TGFβ/BMP signaling is known to play key roles both in the regulation of both tumor growth and osteo-chondrogenic differentiation ^48, 49^ and was found to be active in chondrosarcoma ^50^. Notably, enasidenib treatment induced the downregulation of many of the main factors controlling these signaling pathways. Thus, the expression of ligands such as *TGFB1*, *TGFB2*, *BMP2*, *BMP4* or *BMP7*; receptors like *TFGBR2*, *ACVR1 (ALK2)*, *ACVRL1 (ALK1)* or *BMPR2*; and downstream transcriptional regulators as *SMAD3*, *SMAD5* or *SMAD6* were decreased upon drug treatment. Also, highlighting the complex regulation of this pathway, other important factors in this pathway, such as *BMP6*, *TGFBR1*, *SMAD1* and *SMAD2* were upregulated by enasidenib (Figure 5E). In addition, E2F is a pivotal transcription factor that controls the progression through the cell cycle and its aberrant activation is often responsible for uncontrolled proliferation in cancer cells ^51^. Relevantly, enasidenib induced a broad downregulation of E2F downstream targets in T-CDS17#1 cells, suggesting an inhibition of pathways leading to the activation of E2F-mediated transcription (Figure 5F).

Enasidenib treatment also induced profound deregulation of genes involved in the chondrogenic differentiation process (figure 6A). Many of these genes were downregulated, including those related to TGFβ signaling and other osteochondral differentiation-related genes such as *COL2A1*, *RUNX3*, *ACAN*, *MMP13* or *SOX6*. In any case, other key factors involved in the terminal differentiation of chondrocytes such as *RUNX2* and *SPP1* ^52^ increased their expression following enasidenib treatment.

**Figure 6.**
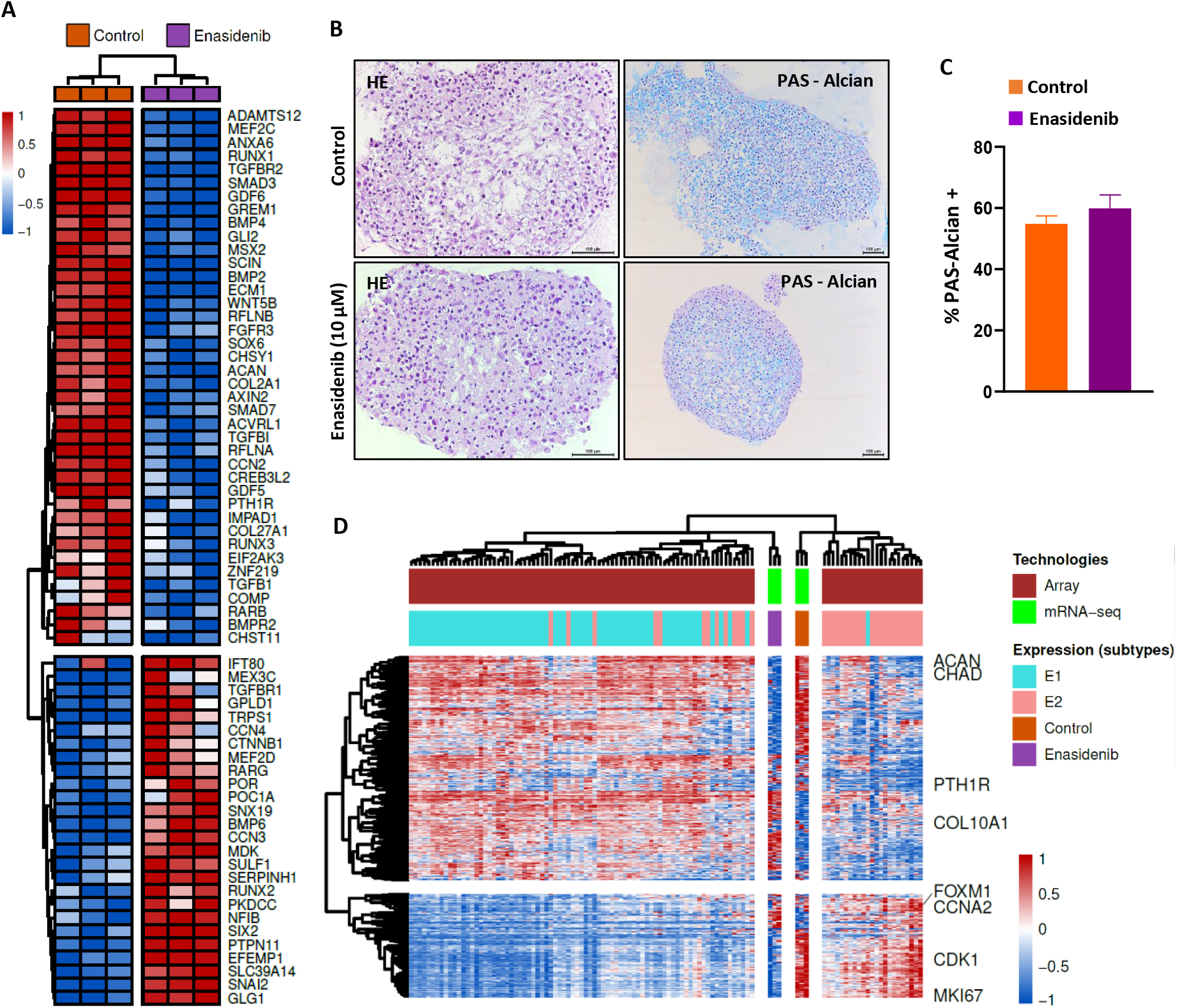
Effect of enasidenib treatment in the chondrogenic differentiation pathway and proliferative status. (A) Heatmap plot depicting the expression values of those DEGs (FDR < 0.05) that belong to the GO BP chondrocyte differentiation pathway. (B-C) Histological analysis of formalin-fixed paraffin-embedded T-CDS17#1 cell spheroids growth in chondrocyte differentiation medium with or without (control) 10 μM enasidenib for 21 days. Representative images of H&E and PAS-alcian staining (B) and quantification of PAS-alcian stain (C) are displayed. Scale bars = 100 μm. Error bars represent the standard deviation of three independent experiments. (D) Heatmap plot showing the expression values of those genes belonging to the mRNA expression signature developed by Nicolle et al. (2019), including their microarray experiments and the classification of chondrosarcoma samples into E1/E2 subtypes.

To check whether these transcriptional enasidenib-induced changes resulted in a functional alteration of the chondrogenic differentiation potential of T-CDS17#1 cells, we performed a chondrogenic differentiation assay in which cell spheroids cultured in differentiation medium were exposed or not to a sub-lethal concentration of enasidenib for 21 days. Despite the transcriptional regulation induced by enasidenib, these experiments showed no differences in the chondrogenic differentiation capability of control and enasidenib-treated T-CDS17#1 cells (Figure 6B-C).

A recent multi-omic analysis of a collection of chondrosarcomas has established a transcriptomic classification that identifies two tumor subtypes (E1 and E2) defined by a balance between chondrogenic tumor differentiation and cell cycle activation ^53^. The E1 phenotype is characterized by the overexpression of chondrogenic differentiation markers such as ACAN, CHAD or PTH1R, while the E2 phenotype is characterized by the activation of proliferative pathways and greater chromosomal instability and is associated with a worse prognosis ^53^. The comparison of the transcriptome of the T-CDS-17#1 cell line, treated or not with enasidenib, with those of the collection of chondrosarcomas described in this work, suggests that treatment with enasidenib can reprogram the transcriptome of our model from a proliferative E2-like phenotype to a less aggressive E1-like one (Figure 6D). In this regard, the repression of the TFGβ pathway and E2F-mediated signaling may underlie the anti-proliferative effect of enasidenib.

### In vivo antitumor activity of enasidenib

Finally, we aimed to test the effectiveness of enasidenib in the in vivo setting. Therefore, we treated immunodeficient mice carrying T-CDS17#1 xenografts with vehicle or 35mg/Kg enasidenib b.i.d. for 21 days. A similar protocol was successfully used for treating in vivo models of AML ^26^. Enasidenib treatment completely inhibited tumor growth and even caused tumor regressions at the experimental end point (Figure 7A). Likewise, tumor weights in the control series doubled those of the enasidenib-treated series at the end of the experiment (Figure 7B). Notably, enasidenib treatment did not cause loss of weight (Figure 7C) or other adverse effects. Histological examination showed that, in agreement with the in vitro assays, enasidenib treatment does not have a relevant impact on the differentiation status of tumors. Thus, both control and enasidenib-treated tumors displayed areas of chondrogenic differentiated (PAS-Alcian positive) and undifferentiated cells (Figure 7D) with not significant changes induced by enasidenib (Figure 7E). On the other hand, enasidenib-treated tumors exhibited a significantly lower proportion of cells staining positive for the proliferation marker ki67 (Figure 7F-G).

**Figure 7.**
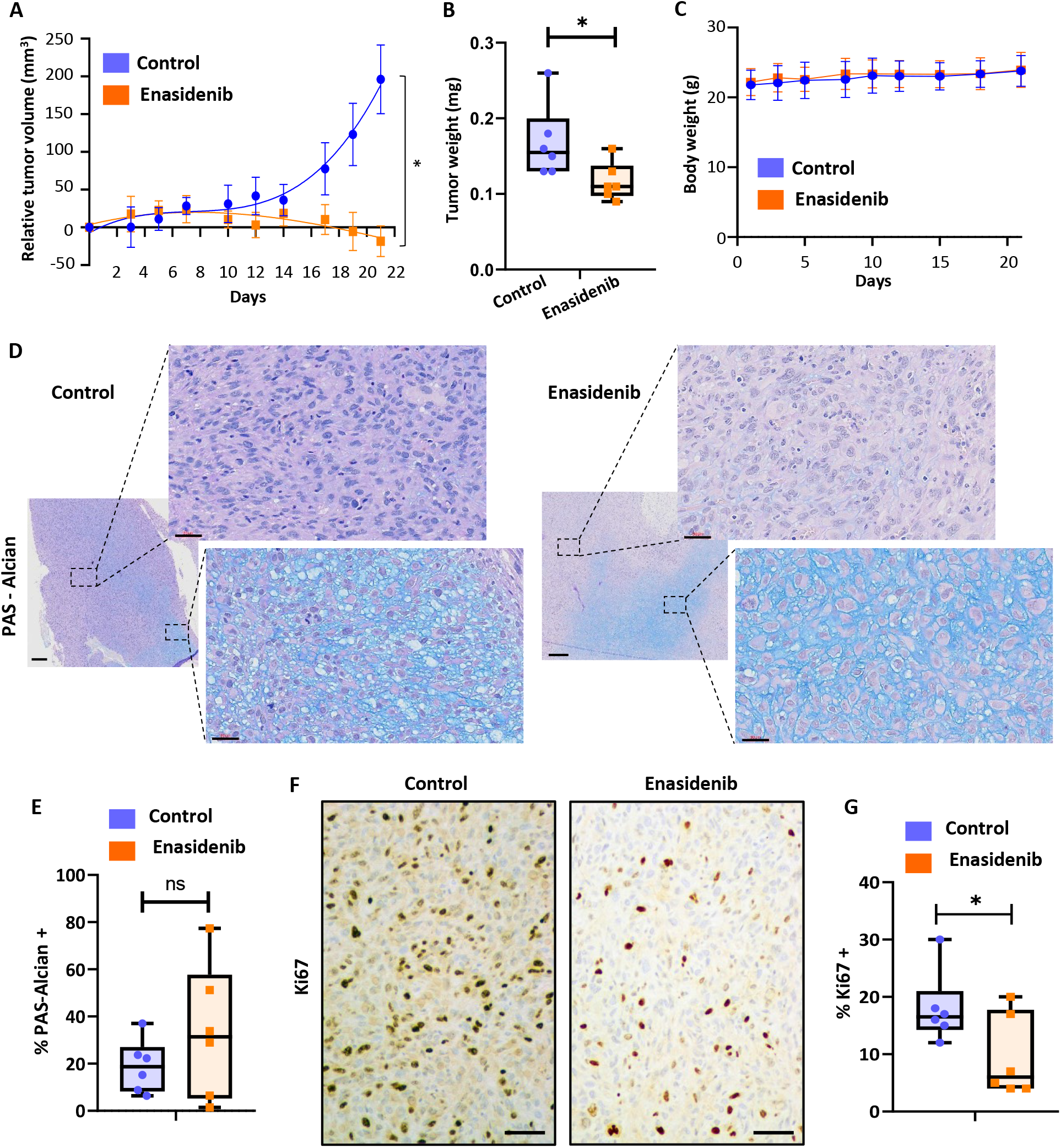
In vivo effect of enasidenib. Established T-CDS17 xenografts were randomly assigned to two different groups (n = 6 per group) and treated b.i.d. with vehicle solvent (control) or 35 mg/kg enasidenib for 21 days. (A) Curves representing the mean relative tumor volumen of T-CDS17 xenografts during the treatments. (B) Average tumor weight at the end of the experiment. (C) Change in the body weights of mice during the treatments. (D-G) Histological analysis of formalin-fixed paraffin-embedded tumors extracted at the end of the treatment. (D) Representative images of PAS-alcian staining of control and treated tumors. Two different areas of the tissue are magnified. Scale bars= 300 µm for left panels and 30 µm for magnified right panels. (E) Quantification of the percentage of PAS-alcian stained areas. (F) immuno-staining detection of Ki67. Scale bars= 100 µm (G) Quantification of Ki67 positive cells. Error bars represent the SD and asterisks indicate statistically significant differences between groups in a two-tailed unpaired t-test (*: p < 0.05; ns: not significant).

Altogether, these results suggest that enasidenib may represent an efficient therapeutic alternative for the case CDS17 and possibly for other chondrosarcomas carrying R172G and R172S variants in the IDH2 gene.

## DISCUSSION

In this work, we have applied NGS protocols to detect targetable mutations in a collection of sarcomas with available patient-derived models. In these analyses, we have detected the oncogenic IDH2-R172G and IDH1-R132L variants in the CDS17 and CDS11 chondrosarcoma models, respectively. These patient-derived cell lines constitute the only models reported to date with these pro-tumor variants in chondrosarcoma. Following this personalized medicine approach, we focus on the specific targeting of IDH mutations in the corresponding patient-derived avatars. First, we analyze for the first time the effect of treating chondrosarcomas with IDH2 mutant inhibitors. We found that the CDS17 / T-CDS17 model carrying the R172G-IDH2 variant and the SW1353 cell line mutated in R172S-IDH2 were much more sensitive to antiproliferative effects of enasidenib than the rest of the cell lines with other mutations in *IDH1/2* or without mutation in these genes. Besides, in vivo treatment with enasidenib of T-CDS17 xenograft models efficiently inhibited tumor growth at non-toxic concentrations. In accordance with the anti-tumor effects produced by enasidenib, we observed that this inhibitor drastically decreased the intracellular levels of D2HG, as previously reported in AML patients treated with enasidenib ^26, 54^.

In contrast to the positive effects of enasidenib, we did not observe any antiproliferative effect when treating our models either with the dual mIDH1/IDH2 inhibitor vorasidenib or with inhibitors of glutaminolysis. A previous study showed that SW1353 and L2975 mIDH2 cell lines respectively highly and moderately sensitive to the in vitro treatment with the glutaminolysis inhibitor CB-839 ^47^. The fact that, unlike the study by Pertese et al, we performed the CB-839 treatments in the presence of serum and using different models of chondrosarcoma may contribute to the different results observed in both studies.

Regarding the specific targeting of mIDH1, previous studies have yielded contradictory results about the anti-tumor potential of mIDH1 inhibitors in chondrosarcomas. On the one hand, one study reported that the mIDH1 inhibitor DS-1001b was capable of inhibiting the in vitro and in vivo proliferation of chondrosarcomas with these mutations ^28^. Likewise, other study conducted with the AGI-5198 inhibitor reported in vitro anti-tumor activity of this drug in IDH1 mutant chondrosarcoma cell lines ^27^. On the other hand, a third study performed in a collection of wild-type and IDH mutant chondrosarcoma cell lines concluded that, despite being able to efficiently decrease D2HG levels, the AGI-5198 inhibitor did not affect the viability, proliferation, migration or methylation status of chondrosarcoma cells ^29^. in any case, Li et al. only observed a cytotoxic effect at the highest concentration assessed which has not been tested by Suijker et al. In line with this last study, we did not observe any effect of ivosidenib on the viability of our panel of chondrosarcoma cell lines, independently of the *IDH1* mutation status.

The most described mechanism of action for inhibitors of mutant IDH enzymes in AML and glioma involves the recovery of the D2HG-inhibited activities of α-KG-dependent dioxygenases and the consequent restoration of normal levels of DNA and histones methylation. This reversal of the hypermethylated phenotype would lead to an increase in differentiation and therefore a decrease in tumor aggressiveness ^23, 24^. However, it has also been reported that not all epigenetic effects caused by *IDH* mutants are reversible ^24, 55^. In addition, clinical responses of AML patients were not always associated with a complete reversal of mutant *IDH*-induced DNA hypermethylation ^24, 56^. In line with these non-canonical cases, we have not found changes in the methylation levels of the CDS17 chondrosarcoma models after treatment with enasidenib. The relevance of IDH mutant-induced epigenetic changes in chondrosarcoma has been revealed in previous studies showing a global hypermethylated status of IDH mutant cases ^53, 57^. Although there could be technical biases arising from the comparison of epigenetic features in tissue samples and cultured cell lines, our methylome analysis of normal cartilage samples and chondrosarcoma cell lines also showed a high rate of hypermethylated genes in IDH2 mutated cells. Moreover, recent work with IDH mutant enchondroma and chondrosarcoma samples indicates that epigenetic remodeling plays an important role in chondrosarcoma progression, although no vulnerabilities specifically associated with IDH mutations could be identified in an epigenetic compound screen ^58^. Altogether, it can be speculated that the intense epigenetic modulation induced by the mutation in our model of dedifferentiated chondrosarcoma has reached a state that cannot be reversed by lowering D2HG levels. In line with this hypothesis, our in vitro and in vivo functional studies showed that enasidenib treatment did not result in increased chondrogenic differentiation, and therefore the observed anti-tumor effects must be mediated by alternative mechanisms. In this sense, the mIDH1 inhibitor DS-1001b was able to induce chondrogenic differentiation in a model of conventional chondrosarcoma but not in a dedifferentiated chondrosarcoma line. Instead, the antitumor mechanism in this second model was associated with the induction of cell cycle arrest ^28^.

Our transcriptomic studies support the involvement of pathways related to the control of cell differentiation and cell fate within the mechanism of action of enasidenib. Among them, signaling mediated by TGFβ and BMPs are among the most strongly repressed by the inhibitor. In chondrosarcoma, these signaling pathways have been reported to be functionally active and are involved in tumor progression and regulation of the dedifferentiated phenotype in high-grade tumors ^50^. We can speculate that the TGFβ/BMP pathway was overactivated in our dedifferentiated chondrosarcoma model and the treatment with enasidenib inhibited this signaling as part of its anti-tumor mechanisms. In addition, our transcriptome analysis provides clues indicating that the anti-tumor effect of enasidenib highly relies on the block of proliferation-related pathways. Thus, within the two types of transcriptomic profiles identified in a multiomic study of a collection of chondrosarcomas ^53^, the transcriptome of T-CDS-17#1 cells corresponded to a proliferative profile associated with a worse prognosis (E2 profile), and the treatment with enasidenib was able to reprogram their transcriptomic landscape towards a phenotype with less aggressive characteristics and better prognosis (E1 profile). Considering the genes of the E1/E2 signatures, we can hypothesize that the inhibitor produces a negative regulation of those genes linked to the cell cycle, proliferation and division, which would explain the general repression of E2F-controlled targets in T-CDS17#1 cells. Relevant to this finding, a recent study found that mutant *IDH*-induced methylation profiles in dedifferentiated chondrosarcomas were associated with upregulated expression of genes involved in G2/M checkpoints and E2F targets ^57^. Among the E2F targets that we found repressed by enasidenib there are key mediators of progression through the G2/M phase of the cell cycle such as *CDK1*, *AURKA*, *PLK1*, *WEE1* or *CHK1*, while the expression of other inhibitors of proliferation such as *TP53*, *CDKN2A* or *CDKN1B* increases after treatment with the inhibitor. Therefore, we can speculate that this blockade of cell growth pathways induced by enasidenib is at the base of its anti-tumor effect. This hypothesis is supported by our in vivo experiments, in which we did not observe cell differentiation in tumors, but we did observe tumor regression associated with a decrease in cell proliferation. Finally, for in vivo experiments we have tested a dose that has been successfully used in preclinical models of AML to support the initiation of clinical trials and the subsequent approval of the drug for the treatment of this disease ^26^. Therefore, the fact, that this concentration was also effective and safe in the chondrosarcoma models assayed in this work, supports future clinical testing in chondrosarcoma patients.

In summary, this study provides the first preclinical evidence of the effectiveness of targeted inhibition of *IDH2* mutants in chondrosarcoma by enasidenib treatment. Our data suggest that the mechanism of action of this drug is associated with the repression of proliferative pathways and not with the promotion of tumor differentiation. This anti-proliferative mechanism of action may be especially relevant in dedifferentiated chondrosarcomas where reversal of this phenotype is not possible. In any case, future research should be designed to define reliable biomarkers capable of identifying those patients who are suitable to benefit from this treatment. Finally, this work provides support for the use of bone sarcoma patient-derived lines as avatar models capable of predicting (pre)-clinical responses in personalized medicine strategies.

## CONTRIBUTORS

**Verónica Rey:** investigation; methodology; data curation; formal analysis; writing—original draft. **Juan Tornín:** investigation; methodology; data curation; formal analysis. **Juan Jose Alba-Linares:** data curation; software; formal analysis; methodology; writing—original draft. **Cristina Robledo:** investigation; methodology; formal analysis. **Dzohara Murillo, Aida Rodríguez, Borja Gallego, and Carmen Huergo:** investigation; methodology. **Alejandro Braña:** resources; methodology. **Aurora Astudillo:** pathological analysis, methodology. **Dominique Heymann:** resources; writing—review and editing. **Agustín Fernández and Mario F. Fraga:** data curation; software; formal analysis; writing—review and editing. **Javier Alonso:** methodology; formal analysis, funding acquisition, writing—review and editing. **René Rodríguez:** Conceptualization; resources; data curation; formal analysis; supervision; funding acquisition; project administration; writing—original draft; writing— review and editing.

## ACKNOWLEDGEMENTS

We acknowledge Daniela Corte-Torres from the Principado de Asturias BioBank (PT17/0015/0023) for its support with histological analysis. In addition, we acknowledge Prof. Judith V.M.G. Bovée and Prof. Karoly Szuhai, both from Leiden University Medical Center, for providing the L2975 cell line and reviewing the manuscript. This work was supported by the Agencia Estatal de Investigación (AEI) [MICINN/Fondo Europeo de Desarrollo Regional (FEDER) (grant PID2019-106666RB-I00 to R.R.) and ISC III/FEDER (grants PI20CIII/00020, DTS18CIII/00005 to J.A.; Consorcio CIBERONC - CB16/12/00390 and Consorcio CIBERER - CB06/07/1009 and CB19/07/00057)]; the Spanish Group for Research on Sarcomas (GEIS) (grant GEIS-62); the Plan de Ciencia Tecnología e Innovación del Principado de Asturias/FEDER [grant IDI/2021/000027 to R.R. and Severo Ochoa predoctoral fellowships BP-20-046 to B.G. and BP-21–084 to DM]; Fundación Científca de la Asociación Española Contra el Cáncer (AECC) (predoctoral fellowship to CH); grant from the Asociación GALBAN-Niños con Cancer to J.T. and R.R.; Asociación Pablo Ugarte (grants TRPV205/18 and DGDO 195/22 to J.A.; Asociación Candela Riera, Asociación Todos Somos Iván and Fundación Sonrisa de Alex (grant numbers TVP333-19, TVP-1324/15) to J.A.

## DATA SHARING

All relevant aggregate data supporting the findings of this study are available within the article and its supplementary information files. Raw data files obtained from RNA seq and DNA methylation analysis have been deposited at the GEO-NCBI repository with SuperSeries reference GSE235701 (https://www.ncbi.nlm.nih.gov/geo/query/acc.cgi?acc=GSE235605 and https://www.ncbi.nlm.nih.gov/geo/query/acc.cgi?acc=GSE235697). Other unprocessed data are available from the corresponding author on reasonable request.

