## Supplemental information for "A personalized medicine approach identifies enasidenib as an efficient treatment for IDH2 mutant chondrosarcoma"

### SUPPLEMENTAL MATERIALS AND METHODS

#### Cell lines

The identity of cell lines has been authenticated by Short Tandem Repeats analysis. All cultures were regularly tested and found to be negative for mycoplasma contamination using the Biotools Mycoplasma Gel Detection kit (B&M LABS, Madrid, Spain).

**Next Generation Sequencing**

*library preparation*

All steps of library preparation were performed according to the manufacturer's protocol (see Supplemental information). Briefly, the amplicons were pooled, purified, labeled with a compatible sequencing adapter/index using NEBNext kit (New England BioLabs, Evry, France) and finally purified again. The quality of the libraries was confirmed using the Agilent high-sensitivity DNA kit (Agilent Technologies, Santa Clara, CA, USA) to confirm the successful generation of 300-bp products for the GeneRead Human Comprehensive Cancer Panel. The individual libraries for each patient were quantified with Kapa Library Quantification Kit (Kapa Biosystems, Manufacturing, R&D, Cape Town, South Africa).

*Identification of pathogenic mutations*

For the identification of somatic mutations, tumor and matched reference were compared between them using the Analyze GeneGlobe (https://geneglobe.qiagen.com/es/) software to detect single nucleotide variants (SNVs), short insertions and deletions (indels) and copy-number variants (CNVs). Variants categorized as non-synonymous, frameshift, stop gain or affect a conserved splice-site in any of the annotated transcripts associated with the cancer gene. The variants were classified as pathogenic or probably pathogenic. The resulting somatic variants with more than 100 reads coverage and a VAF above 5% were manually reviewed with the IGV browser (Integrative Genomics Viewer, Broad Institute, CA, USA) to detect possible sequencing artefacts, for example in poorly mapped regions, last base of amplicons or polynucleotide tracks. Variants that were observed in the control sample and suspected of being sequencing errors by visual inspection on the IGV browser were filtered-out.

**RNA sequencing**

*RNA-seq data preprocessing:* All sequencing data files were evaluated using the *fastQC* software (v.0.11.9) for quality control. Adapters and poor-quality paired-end reads were removed using *fastp* (v.0.20.1) with the following options: --detect_adapter_for_pe --trim_poly_x --correction -r -M 10 -l 20.

*Transcript-level quantification of RNA-seq data:* preprocessed reads were pseudo-aligned to the *GRCh38* human genome and quantified at the transcript-level using the mapping-based mode of the *Salmon* package (v.1.3.0) with the following options: --validateMappings --gcBias. A *Salmon* index was previously generated from a *gentrome* file which was constructed by the *gencode.v36* transcriptome and the *GRCh38.p13* DNA primary assembly.

*Differential gene expression analyses:* Analyses of gene expression data were performed using R statistical software (v.4.0.2). Transcript-level information was summarized at the gene-level for both exploratory and differential analyses using the tximport package (v.1.16.1). For visualization purposes, the matrix of raw counts was *rlog* (regularized logarithm) transformed according to the *DESeq2* package (v.1.28.1). Not or low-expressed genes (< 2 counts across all conditions) were discarded to reduce the multiple testing penalty. Differential gene expression analyses were then performed using the negative binomial generalized linear model fitting implemented in *DESeq2*, with counts normalized by the mean ratio method, which corrects for sequencing depth and RNA composition biases. After adjusting *p*-values for multiple testing using the Benjamini-Hochberg method, differentially expressed genes (DEGs) were defined as those with FDR<0.05.

*Gene annotation:* The *org.Hs.eg.db* package (v.3.11.4) was used to annotate each Ensembl Stable Gene ID with their corresponding Entrez Gene ID and Gene Symbol categories.

*Gene set enrichment analyses:* To interrogate the functionality of the DEGs, enrichment analyses were performed with the *fgsea* package (v.1.14.0) on the gene sets from the *Molecular Signatures Database* (MSigDB), accessed via the *msigdbr* package (v.7.2.1). Gene ranking was provided to the *fgsea* algorithm using the log-transformation of *p*-values [$-\log_{10}(p\text{-values}) \; x \; sign(\log_2 FC)$, FC: fold change)].

*Microarray analysis of chondrosarcoma samples:* Microarray expression data (Affymetrix Human Gene 2.0 ST arrays) from Nicolle et al. (2019) were analyzed to determine whether enasidenib treatment can alter transcriptome profiles associated with chondrosarcoma expression subtypes. Log expression values from microarray and mRNA-seq experiments were independently normalized to the [0,1] range to reduce bias due to the different expression technologies considered, and samples were hierarchically clustered using Euclidean distances.

**Genome-wide DNA methylation analysis**

*Infinium MethylationEPIC data preprocessing:* MethylationEPIC BeadChip data analyses were performed using the *minfi* package (v.1.32.0) in statistical software R (v.4.0.2). All samples validated the expected sex identity of the cell line, preserved the methylation landscape of SNP probes and passed the specific quality control for intensity signals in both methylated and unmethylated channels. After checking the quality of the samples, the ssNoob method was applied to correct background noise signals affecting methylation intensity values. β-values were calculated for each interrogated CpG site using the following formula: $\beta = \frac{max(y[meth],0)}{max(y[meth],0)+max(y[unmeth],0)+100}$. Next, the beta-mixture quartile normalization (BMIQ) implemented in *ChAMP* (v.2.16.2) was selected to overcome the probe design bias of Illumina Infinium arrays. Finally, to avoid spurious methylation signals due to technical issues or unwanted biological sources of variation, probes were filtered according to the following exclusion criteria: (a) detection p-value > 0.01 in any sample; (b) cross-reactive or multi-mapping nature, (c) sex chromosome location and (d) inclusion of SNPs with MAF ≥ 0.01 at their CpG or SBE sites (dbSNP v.147).

*Differential methylation analyses:* Linear regression models were built to detect differentially methylated probes (DMPs) using the *limma* package (v.3.44.3), with the methylation value as the dependent variable. To increase homoscedasticity in the models, β-values were logit-transformed to M-values using the beta2m function of the lumi package (v.2.40.0). Sex and study accession code were included as covariates in the linear modelling. Empirical Bayes-moderated t-tests were employed to define contrasts and the p-values were adjusted for multiple comparisons using the Benjamini-Hochberg method (FDR<0.05). In addition, filtering based on the absolute difference in beta values (|Δβ|) for each pairwise comparison (>20%) was applied to define functionally relevant DMPs. To detect differentially methylated regions (DMRs), the limma p-values from DMPs were inputted in the comb-p function of the Enmix package (v.1.28.2) using default parameters, discovering CpG sites that were spatially related at the level of statistical significance. DMRs were then selected under the following thresholds: (a) FDR<0.05, (b) Sidak-corrected p-value <0.05, (c) $|\Delta\bar{\beta}|$>20%, $\bar{\beta}$: mean methylation value (region). In addition, DMRs displaying less than 66% of CpG sites with changes in the same direction (hyper: > 5 % gain; hypo: > 5 % loss; equal: ≤ 5% change) were filtered out, as were DMRs with less than 5 CpGs and those regions that were not annotated to any gene. Finally, only those genes whose DMRs changed in the same direction were retained as differentially methylated genes.

*Probe annotation:* The *IlluminaHumanMethylationEPICanno.ilm10b4.hg19* package (v.0.6.0) was used to assign each probe to its CGI (CpG Island) and gene location status. For the annotation of regions, the probes belonging to each region were first individually annotated as described above. A single annotation was then assigned to each region according to the following criteria: (1) for CGI status, "Island">"N_Shore">"S_Shore">"N_Shelf">"S_Shelf">"OpenSea"; and (2) for gene locations, "TSS1500">"TSS200">"5'UTR">"1stExon">"Body">"ExonBnd">"3'UTR">"Intergenic".

*Pathway enrichment analyses:* To interrogate the functionality of DMPs, pathway enrichment analyses were performed using the gsameth function of the missMethyl package (v.1.22.0) on the gene sets from the Molecular Signatures Database (MSigDB), accessed via the msigdbr package (v.7.2.1). The number of probes mapping to each gene was taken into account as a bias factor for the enrichment analyses.

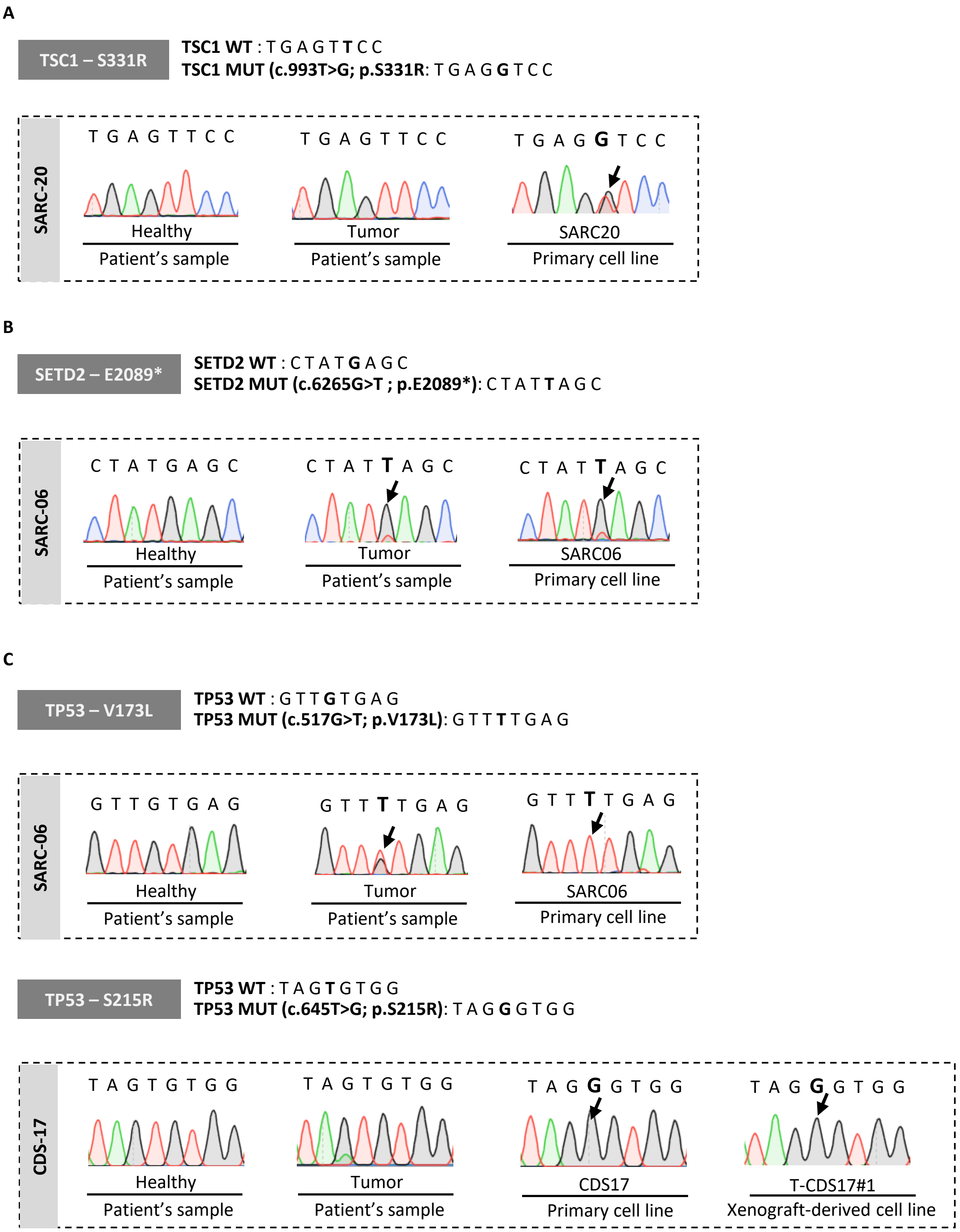


**Figure S1. Sanger sequencing validation of mutations detected in TSC1, SETD2 and TP53 genes.** Sanger sequencing chromatograms showing mutations (black arrows) detected in TSC1 (A), SETD2 (B) and TP53 (C) genes present in the indicated healthy tissue, tumor tissue and cell lines. Reference wild type (WT) and mutated (MUT) sequences for each gene are shown.

**Rey et al. Figure S2**

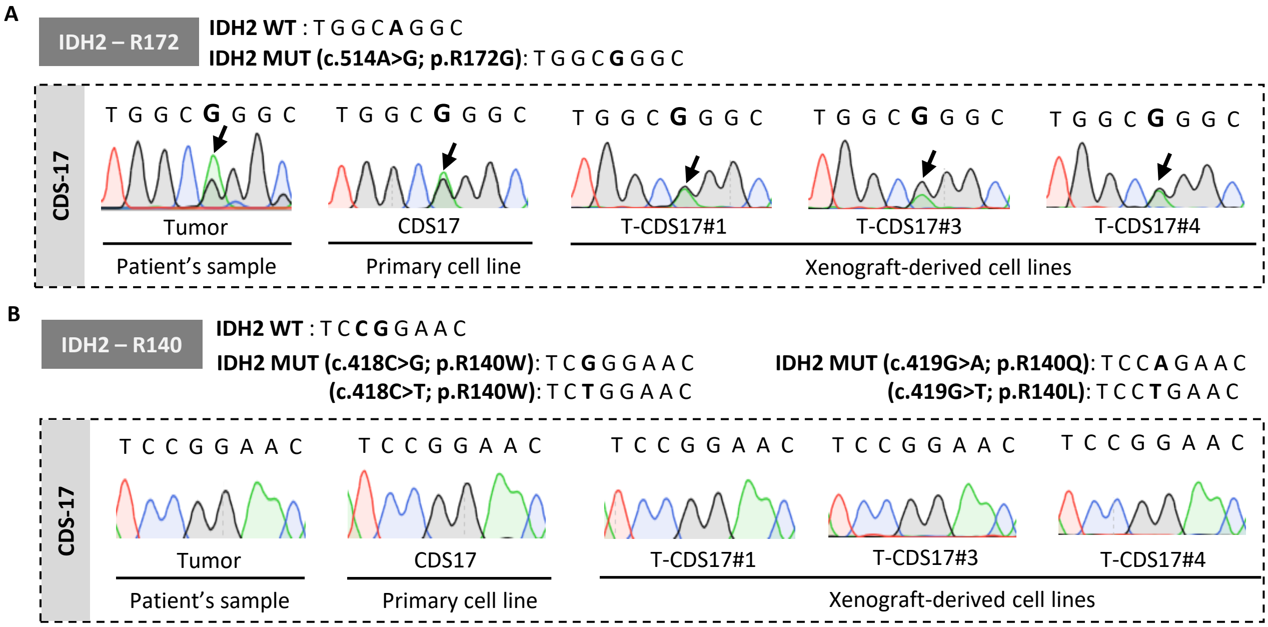


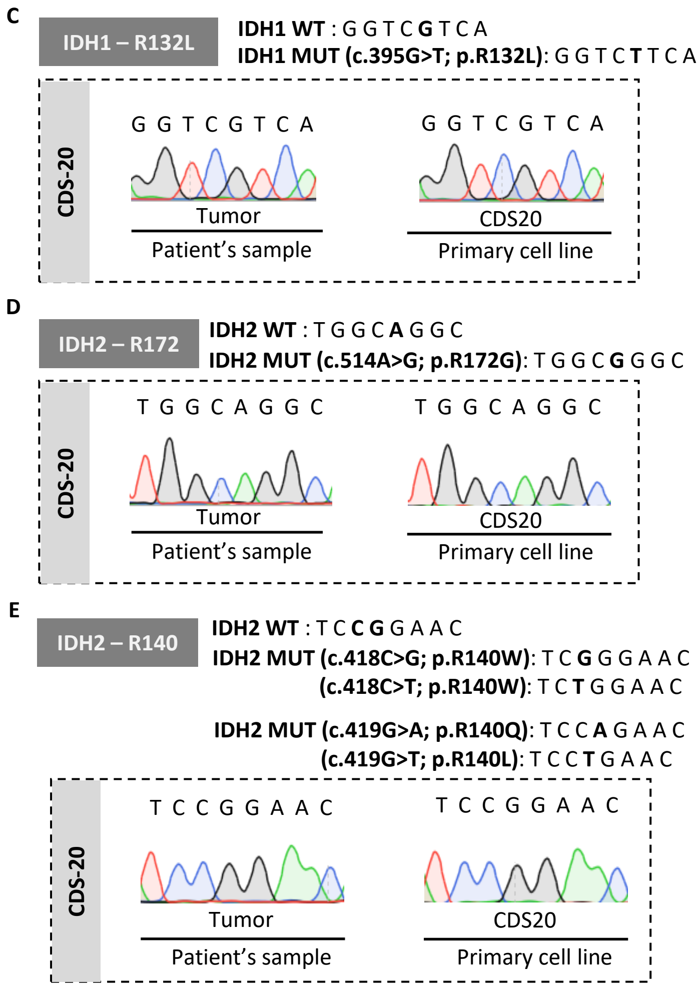


**Figure S2. . Sanger sequencing of IDH1 and IDH2 genes in CDS17 and CDS20 models.** Sanger sequencing of mutation hotspots in the IDH2 gene in tumor tissue, primary cell line, and cell lines derived from xenografts of the CDS17 model. Chromatograms show non-synonymous mutations (black arrows) at residue R172 (A) and wild type sequence at residue R140 (B). (C-E) Sequencing of tumor tissue and primary cell line of the CDS20 model showing no motutations in IDH1-R132L (C), IDH2-R172 (D) and IDH2-R140 (E) hotspots.

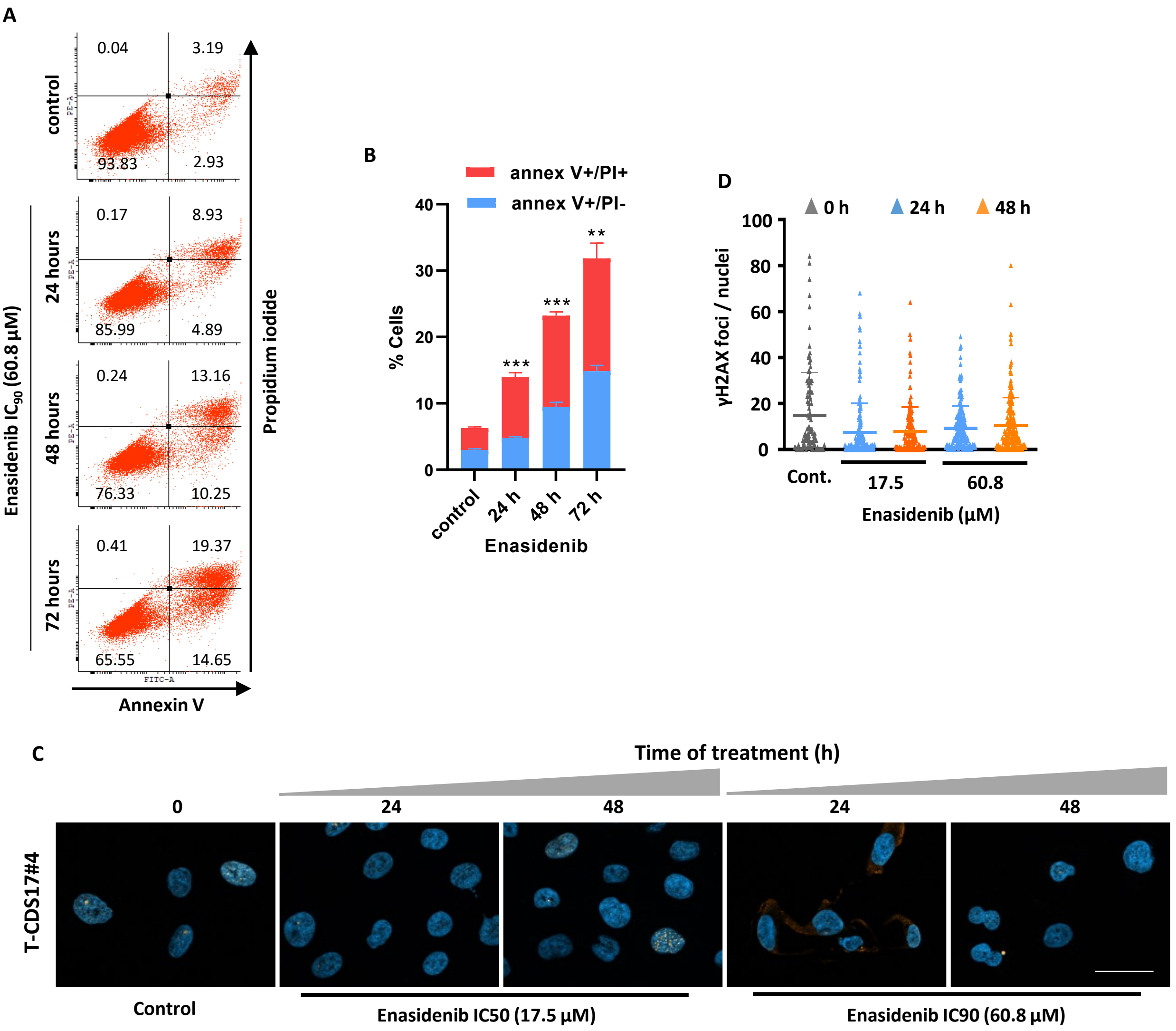


**Figure S3. Analysis of apoptosis and DNA damage in enasidenib treated chondrosarcoma cells.** (A-B) Annexin V / propidium iodide binding assay of T-CDS17#4 cells treated with 60.8 µM enasidenib for 24, 48 and 72 hours. Representative dot plots (A) and summary (mean and SD) of three independent experiments (B) are shown. Asterisks indicate statistically significant differences in the values of accumulated Annexin V+ cells with the control series (**: $p < 0.01$; ***:$p < 0.001$, one-way ANOVA). (C-D) Analysis of γ-H2AX foci formation after treatment of T-CDS-17#4 cells with $IC_{50}$ (17.5 µM) and $IC_{90}$ (60.8 µM) concentrations of enasidenib for 0 h (control), 24 h and 48 h. (C) Representative images of immunostaining experiments (γ-H2AX immunodetection: red fluorescence; DAPI staining: blue fluorescence) for each condition. Scale bars = 25 µm. (D) Quantification of γ-H2AX foci. Means (horizontal bars) and SD of the number of foci of at least 100 cells for each condition are shown.

**Table S1.** Patient and tumor characteristics

| Cell line | Xenograft line | Age* | Gender | Tumor location | Grade | Diagnosis |
|---|---|---|---|---|---|---|
| **CDS11** | - | 45 | female | femur | II | conventional chondrosarcoma |
| **CDS17** | **T-CDS17#1 / #3 / #4** | 49 | male | hemipelvis | III | dedifferentiated chondrosarcoma |
| **CDS20** | **-** | 67 | male | knee | III | extraskeletal myxoid chondrosarcoma |
| **SW1353** | **-** | 72 | female | humerus | II | central chondrosarcoma |
| **L2975** | **-** | 57 | male | femur | High | dedifferentiated chondrosarcoma |
| **SARC06** | - | 52 | male | Thigh | III | undifferentiated pleomorphic sarcoma |
| **SARC20** | - | 57 | male | Thigh | III | undifferentiated pleomorphic sarcoma |
| **SYN01** | - | 24 | male | leg | II | biphasic synovial sarcoma |

(* )Age at diagnostic.

**Table S2. Genes included in the QIAseq Human Comprehensive Cancer Panel.**

| | | | | | | | |
|---|---|---|---|---|---|---|---|
| *ABL1* | *BUB1B* | *DDR2* | *FGFR2* | *IDH2* | *MEN1* | *PDGFRA* | *SMARCA4* |
| *AKT1* | *CARD11* | *DICER1* | *FGFR3* | *IKZF1* | *MET* | *PHF6* | *SMARCB1* |
| *AKT2* | *CBL* | *DNMT3A* | *FH* | *IL6ST* | *MLH1* | *PIK3CA* | *SMO* |
| *ALK* | *CBLB* | *ECT2L* | *FLCN* | *IL7R* | *MSH2* | *PIK3R1* | *SPOP* |
| *AMER1* | *CD79A* | *EGFR* | *FLT3* | *JAK1* | *MSH6* | *PMS2* | *SRC* |
| *APC* | *CD79B* | *EP300* | *FUBP1* | *JAK2* | *MTOR* | *PPP2R1A* | *STK11* |
| *AR* | *CDC73* | *EPCAM* | *GATA1* | *JAK3* | *MUTYH* | *PRDM1* | *SUFU* |
| *ARID1A* | *CDH1* | *ERBB2* | *GATA2* | *KDM6A* | *MYC* | *PRKAR1A* | *TERT* |
| *ARID2* | *CDK12* | *ERBB3* | *GATA3* | *KDR* | *MYD88* | *PTCH1* | *TNFAIP3* |
| *ASXL1* | *CDK4* | *ERBB4* | *GNA11* | *KIT* | *NF1* | *PTEN* | *TNFRSF14* |
| *ATM* | *CDKN2A* | *ERCC5* | *GNAQ* | *KLF6* | *NF2* | *PTPN11* | *TP53* |
| *ATRX* | *CHEK2* | *ESR1* | *GNAS* | *KMT2D* | *NFE2L2* | *RAC1* | *TSC1* |
| *BAP1* | *CIC* | *EZH2* | *GPC3* | *KRAS* | *NFKBIA* | *RB1* | *TSC2* |
| *BCL6* | *CREBBP* | *FAM46C* | *GRIN2A* | *MAP2K1* | *NOTCH1* | *RET* | *TSHR* |
| *BCOR* | *CRLF2* | *FANCA* | *H3F3A* | *MAP2K2* | *NOTCH2* | *ROS1* | *U2AF1* |
| *BRAF* | *CSF1R* | *FANCD2* | *HIST1H3B* | *MAP2K4* | *NPM1* | *SDHB* | *VHL* |
| *BRCA1* | *CTNNB1* | *FANCE* | *HNF1A* | *MAP3K1* | *NRAS* | *SETD2* | *WT1* |
| *BRCA2* | *CYLD* | *FAS* | *HRAS* | *MAP4K3* | *PALB2* | *SF3B1* | *XPC* |
| *BRIP1* | *DAXX* | *FBXO11* | *HSPH1* | *MDM2* | *PAX5* | *SLC7A8* | *ZNF2* |
| *BTK* | *DDB2* | *FBXW7* | *IDH1* | *MED12* | *PBRM1* | *SMAD4* | *ZRSR2* |

**Table S3. Primers for Sanger sequencing**

| GENE | EXON | CODON | PRIMERS | | REFERENCE SEQUENCE |
|---|---|---|---|---|---|
| | | | Fw (5´- 3´) | Rv (5´- 3´) | |
| IDH1 | 4 | 101 to 133 | CACCAAATGGCACCATACGA | CAATTTCATACCTTGCTTAATGGG | NM_005896.4 |
| IDH2 | 4 | 126 to 178 | ATTCTGGTTGAAAGATGGCG | AAGTCTGTGGCCTTGTACTG | NM_002168.2 |
| SETD2 | 14 | 2054 to 2277 | GCGCTCGAAAGAACAAATCT | CTGGGTAGATGACGGAGGAG | NM_001349370.2 |
| TSC1 | 10 | 305 to343 | TATCCATGGCCAGTGTTCAG | CCCTGTGTTCTTCTCTTCCATT | NM_000368.4 |
| TP53 | 5 | 126 to 187 | CAGTTGCAAACCAGACCTCA | GTTTCTTTGCTGCCGTCTTC | NM_000546.5 |
| | 6 | 187 to 224 | GCCTCTGATTCCTCACTGAT | GCCACTGACAACCACCCTTA | NM_001126114 |
